# Dietary sugar inhibits satiation by decreasing the central processing of sweet taste

**DOI:** 10.1101/2019.12.16.877613

**Authors:** Christina E. May, Julia Rosander, Jen Gottfried, Evan Dennis, Monica Dus

## Abstract

From humans to flies, exposure to diets rich in sugar and fat lowers taste sensation, changes food choices, and promotes feeding. However, how these peripheral alterations influence eating is unknown. Here we used the genetically tractable organism D. melanogaster to define the neural mechanisms through which this occurs. We characterized a population of protocerebral anterior medial dopaminergic neurons (PAM DANs) that innervates the β’2 compartment of the mushroom body and responds to sweet taste. In animals fed a high sugar diet, the response of PAM-β’2 to sweet stimuli was reduced and delayed, and sensitive to the strength of the signal transmission out of the sensory neurons. We found that PAM-β’2 DANs activity controls feeding rate and satiation: closed-loop optogenetic activation of β’2 DANs restored normal eating in animals fed high sucrose. These data argue that diet-dependent alterations in taste weaken satiation by impairing the central processing of sensory signals.

## Introduction

Consumption of diets high in sugar and fat decreases the perception of taste stimuli, influencing food preference and promoting food intake (Bartoshuk et al. 2006; Sartor et al. 2011; Kaufman et al. 2018; Ahart et al. 2019; May et al. 2019; Weiss et al. 2019). Recent studies have examined the effects of these diets on the sensitivity of the peripheral taste system and the intensity of taste experience (Maliphol, Garth, and Medler 2013; Kaufman et al. 2018; May et al. 2019; Weiss et al. 2019), but how exactly taste deficits increase feeding behavior is not known. Orosensory signals determine the palatability or “liking” for foods (Berridge and Kringelbach 2015), but they also promote meal termination via a process called “sensory-enhanced (or mediated) satiety” (Chambers, McCrickerd, and Yeomans 2015). Indeed, foods that provide longer and more intense sensory exposure are more satiating, reducing hunger and subsequent test-meal intake in humans (Cecil, Francis, and Read 1998; Bolhuis et al. 2011; Viskaal-van Dongen, Kok, and de Graaf 2011; Yeomans and Chambers 2011; Forde et al. 2013; Ramaekers et al. 2014). Specifically, sensory signals are thought to function early in the satiety cascade (J. E. Blundell, Rogers, and Hill 1987) by promoting satiation and bringing the on-going eating episode to an end (J. Blundell et al. 2010; Bellisle and Blundell 2013). This is in contrast to nutrient-derived signals, which develop more slowly and consolidate satiety by inhibiting further eating after the end of a meal (J. Blundell et al. 2010; Bellisle and Blundell 2013). We reasoned that if orosensory attributes like taste intensity are important to curtail a feeding event, then diet-dependent changes in taste sensation could promote feeding by impairing sensory-enhanced satiation. Here we investigated the relationship between diet composition –specifically high dietary sugar – the central processing of sweet taste signals, and satiation by exploiting the simple taste system and the conserved neurochemistry of the fruit fly *D. melanogaster*.

Like humans and rodents, fruit flies exposed to palatable diets rich in sugar or fat overconsume, gain weight, and become at-risk for obesity and metabolic syndrome (Musselman and Kühnlein 2018). We recently showed that, in addition to promoting feeding by increasing meal size, consumption of high dietary sugar decreased the electrophysiological and calcium responses of the *Gr64f+* sweet sensing neurons to sweet stimuli, independently of weight gain (May et al. 2019). These physiological changes in the *Gr64f+* cells reduced the fruit flies’ taste sensitivity and response intensity. Opto- and neurogenetics manipulations to correct the responses of the *Gr64f+* neurons to sugar prevented animals exposed to high dietary sugar from overfeeding and restored normal meal size (May et al. 2019). Thus, the diet-dependent dulling in sweet taste causes higher feeding in flies, but how does this happen? How do alterations in the peripheral sensory neurons modulate a behavior as complex as feeding? To better understand how this occurs, we decided to examine the effects of high dietary sugar and taste changes in the central processing of sweet stimuli by dopaminergic neurons (DANs) in the Protocerebral Anterior Medial (PAM) cluster, which respond to the sweet sensory properties to signal sugar reward (Burke et al. 2012; Liu et al. 2012) and reinforce short term appetitive memories (Yamagata et al. 2015; Huetteroth et al. 2015). We hypothesized that impairments in the peripheral responses to sugar could influence the way sweet taste information is transduced through PAM-DANs to affect feeding.

We found that in flies fed a high sugar diet the presynaptic responses of a specific subset of PAM DANs to sweet taste are decreased and delayed. These changes are specific to sweet stimuli and mediated by high dietary sugar. Further, we show that the reduction in the central processing of sweet taste information increases the duration and size of meals: closed-loop optogenetic stimulation of a specific set of PAM DANs corrected meal size, duration, and feeding rate. Together, our results argue that diet-dependent alterations in the central processing of sweet sensory responses delay meal termination by impairing the process of sensory-enhanced satiation.

## RESULTS

### Consumption of a high sugar diet decreases and delays the central processing of the sweet taste signal

In the absence of mapped II-order labellar sweet taste neurons, we used the genetically encoded vesicular release sensor *synaptobrevin-pHluorin* (*syb-pHluorin)* (Poskanzer et al. 2003) to ask if the transmission of the sweet taste signal out of the *Gr64f+* sensory neurons was decreased. We measured the *in vivo* fluorescence from the *Gr64f+* presynaptic terminals in the Sub Esophageal Zone (SEZ) in response to 30% sugar stimulation of the proboscis. We found that the *syb-pHluorin* fluorescent changes upon sugar presentation were markedly decreased when flies were fed a high sugar diet (SD, 30% sucrose) for 7 days, compared to age-matched flies fed a control diet (CD, ∼8% sucrose) (Figure 1A, B). These data suggest that both the responses of the sweet sensing *Gr64f+* neurons to sugar and the transmission of the sweet taste signal are impaired by exposure to the SD.

**Figure 1:**
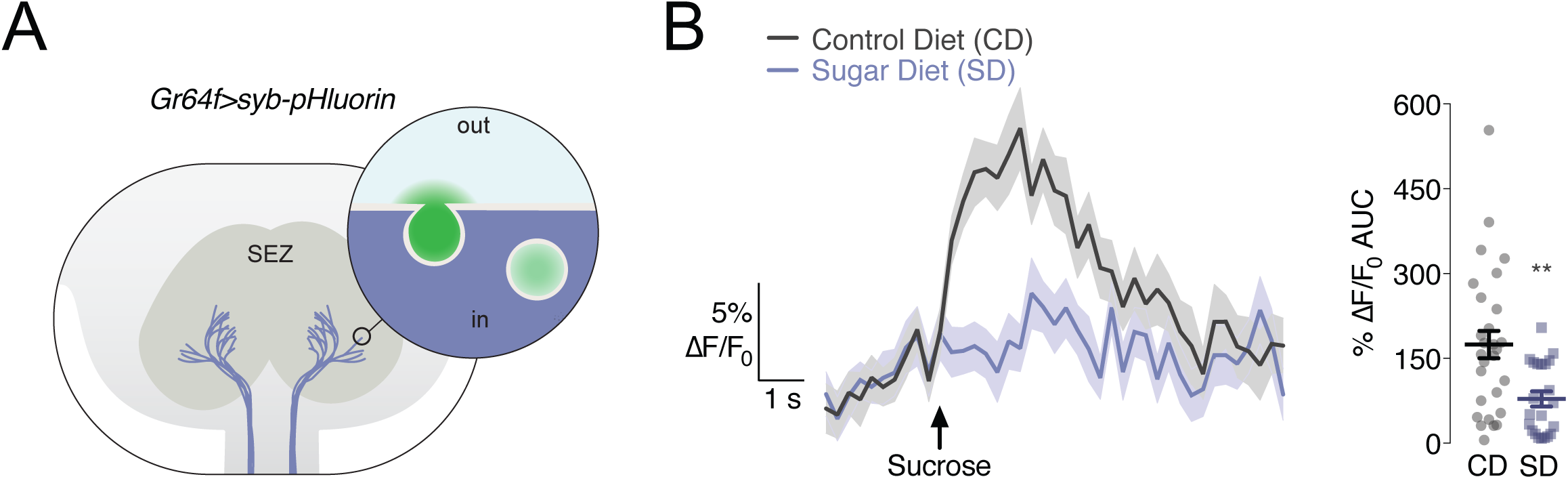
Vesicular release from the *Gr64f+* taste neurons in response to a sucrose stimulus is decreased in flies fed a high sugar diet. A. Schematic of the subesophageal zone (SEZ), highlighting the *Gr64f+* neuron terminals in lavender. Popout bubble demonstrates increased fluorescence upon vesicular release. B. *Left,* Mean %ΔF/F_0_ response traces and *Right*, Area-under-the-curve (AUC) value of %ΔF/F_0_ responses when *Gr64f>syb-pHluorin* flies fed a control (CD, *grey*) or sugar diet (SD, *lavender*) were stimulated with 30% sucrose on the labellum. n=22-28; shading and error bars depict the standard error of the mean. Mann-Whitney test; *** p<0.001.

While the neural pathways that bring sensory information from the periphery to the higher order brain regions are unique across organisms, dopaminergic circuits dedicated to the central processing of sweet taste information exists in humans, rodents, and fruit flies; interestingly, the taste and nutrient properties of sugar are relayed via distinct pathways in these organisms (Yamagata et al. 2015; Huetteroth et al. 2015; Tellez et al. 2016; Thanarajah et al. 2019). Since the involvement of DANs in feeding behavior and in central processing of sensory information is a homologous feature, we decided to center on this circuit as a possible link between diet-dependent changes in sweet responses, higher feeding, and weight gain. In flies, DANs in the Protocerebral Anterior Medial (PAM) cluster that are labeled by the *R48B04-GAL4* transgene and innervate the β’2 and γ4 compartments of the Mushroom Body (MB), respond to sweet sensory properties (Huetteroth et al. 2015; Yamagata et al. 2015); neurons of this population also centrally reinforce water taste (Lin et al. 2014). Here we focused on the β’2 compartment because of its role in processing of the taste properties alone, compared to γ4 which is modulated by both taste and additional factors, such as internal state (Lin et al. 2014; Yamagata et al. 2015). In addition to labeling ∼60 DANs in each PAM cluster, *R48B04* is expressed in other neurons, including in the SEZ. To avoid potential confounding effects of its expression in the SEZ, we used FlyLight to visually identify *GAL4* lines that label subsets of PAM-β’2, but do not label neurons in the SEZ (Aso and Rubin 2016), and identified the split-*GAL4* line *MB301B* which labels ∼12 TH+ PAM-β2β’2a (Figure 2A and Supplemental Figure 1A). We then used the presynaptically targeted *GCaMP6s::Bruchpilot::mCherry* (Kiragasi et al. 2017) to record the response of *MB301B* neurons to stimulation of the labellum with 30% sucrose. We observed an increase in signal in the β’2 compartment, showing that these PAM-β’2 neurons process sweet sensory information (Figure 2B, grey lines). Next we measured the responses of *MB301B* neurons to sucrose taste in flies fed a SD for 7 days and we found a nearly 50% decrease (Figure 2B, rose lines). Furthermore, when we looked at both the average and individual traces, we saw a ∼600 millisecond delay in the peak responses to the sucrose stimulus delivery to the labellum (Figure 2C). No sugar taste responses were recorded in β2, consistent with the idea that it is not involved in taste processing (Figure 2D). Thus, the central processing of sweet stimuli in PAM-β’2 *MB301B* neurons is both decreased and delayed by exposure to a high sugar diet.

**Figure 2:**
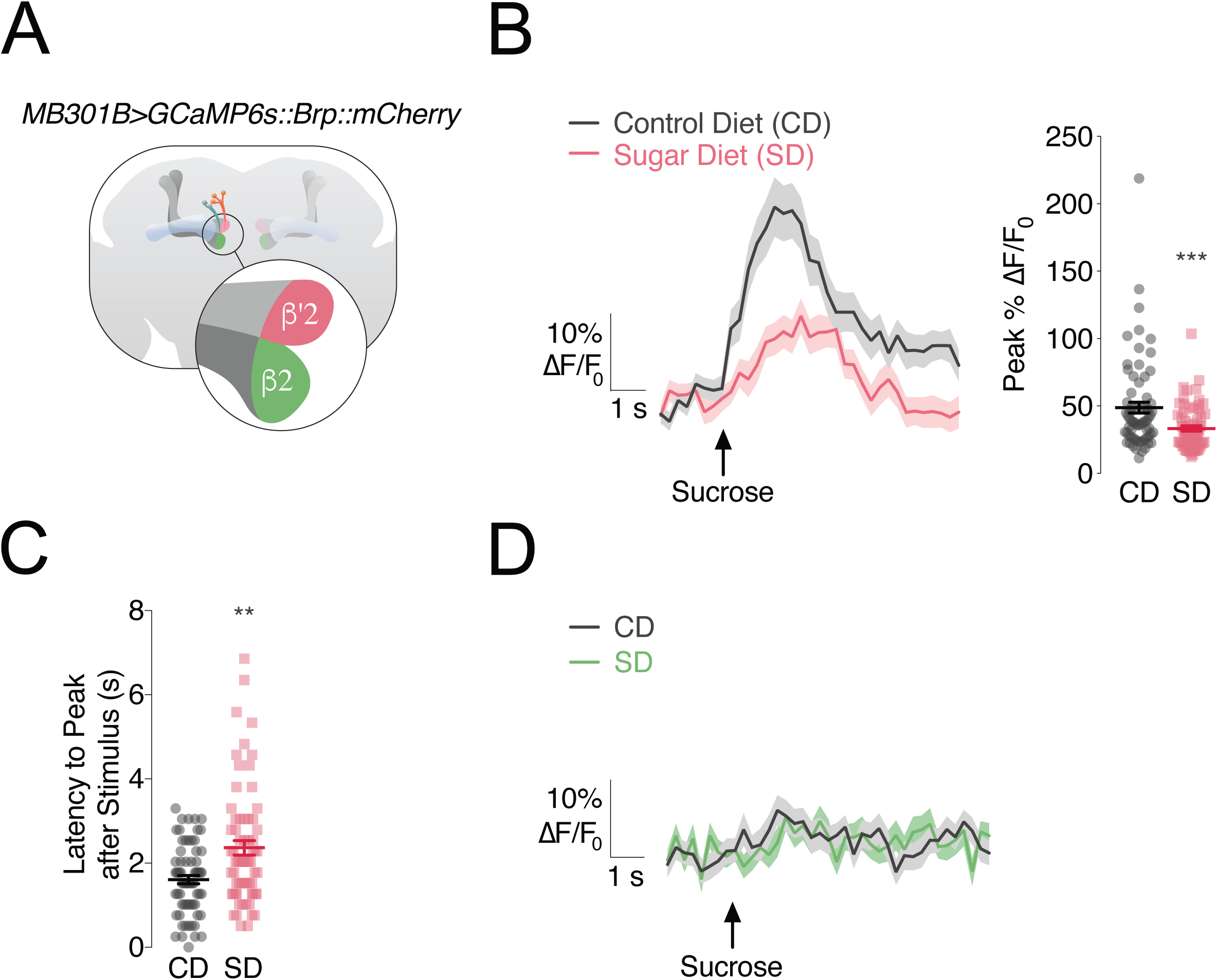
The responses of PAM-β’2 neurons to sweet stimuli change in flies fed a high sugar diet. A. Anatomy of the Mushroom Body (MB) of the *Drosophila melanogaster* brain, with α/β, α’/β’ lobes in greys, γ in translucent blue, and *MB301B* neurons in rose and green; *popout bubble*, schematic showing the β’2 (*rose*) and β2 (*green*) compartments in their respective MB lobes. B. *Left,* Mean %ΔF/F_0_ traces and *Right,* quantification of the maximum peak %ΔF/F_0_ responses to 30% sucrose stimulation of the labellum in the β’2 compartment of *MB301B>GCaMP6s::Brp::mCherry* flies fed a control (CD, *grey*) and sugar diet (SD, *rose*). n=67-70; Shading and error bars are standard error of the mean. n=67-70; Mann-Whitney test; *** p<0.001. C. The delay in the calcium responses quantified as latency in seconds (s) to maximum peak ΔF/F_0_ from the animals in B. n=67-70; Mann-Whitney test; ** p<0.01. D. Mean %ΔF/F_0_ traces for the responses to 30% sucrose stimulation of the labellum in the β2 compartment of *MB301B>GCaMP6s::Brp::mCherry* flies fed a control (CD, *grey*) and sugar diet (SD, *green*). n=67-70; shading is standard error of the mean.

### Alterations in PAM-β’2 responses are specific to high dietary sugar and sweet stimuli

The reduction and delay in central responses to sugar taste in PAM-β’2 DANs on a SD could be due either to the lower transmission of the sensory signal out of the peripheral sweet taste neurons (Figure 1) or to the metabolic side effects of the high nutrient diet. To differentiate between these possibilities, we took multiple approaches. In addition to sweet stimuli, PAM-β’2 neurons also respond to water (Lin et al. 2014); we reasoned that if high dietary sugar unspecifically changed the activity of the PAM-β’2, we would expect flies on the SD to also exhibit impaired central responses to water. However, the magnitude and timing of the β’2 response to water stimulation of the labellum was unchanged between flies on a CD or SD (Figure 3A, B, water stimulation was delivered in the same flies as in Figure 2). Thus, the decrease in PAM-β’2 responses in flies fed a SD is specific to the sweet sensory stimulus. This argues that the overall ability of these DANs to respond to stimuli is not generally affected, and the reduction observed on a SD could occur because of the diet-dependent changes in the sweet taste neurons in the periphery ((May et al. 2019) and Figure 1).

**Figure 3:**
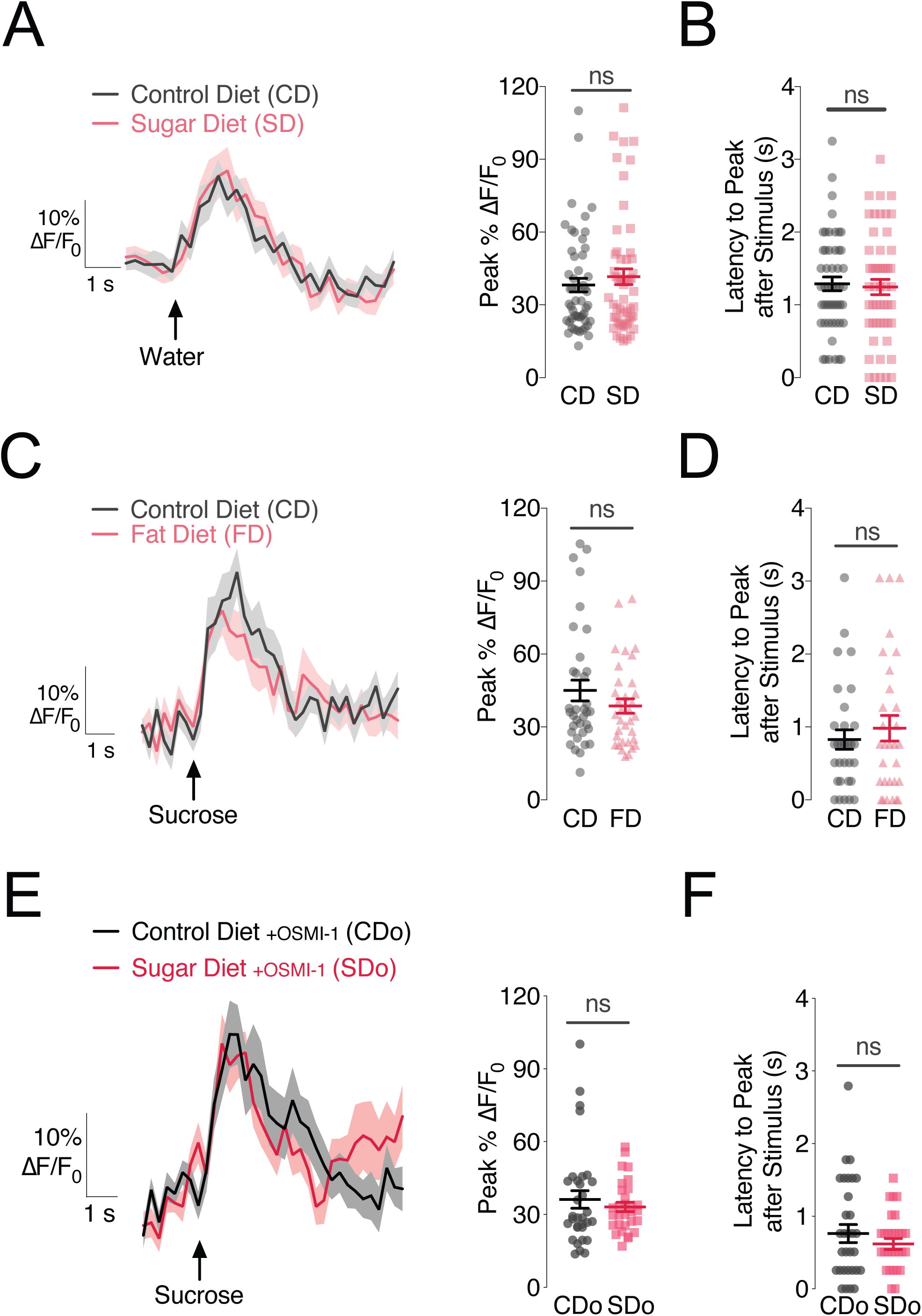
The changes in PAM-β’2 activity are specific to sugar stimuli and caused by deficits in sweet sensation. A. *Left,* Mean %ΔF/F_0_ traces and *Right,* quantification of the maximum peak %ΔF/F_0_ responses to water stimulation of the labellum in the β’2 compartment of *MB301B>GCaMP6s::Brp::mCherry* flies fed a control (CD, *grey*) and sugar diet (SD, *rose*), same animals as in Figure 2. n=67-70; shading and error bars are standard error of the mean. Mann-Whitney test; no significance. B. The delay in the calcium responses quantified as latency in seconds (s) to maximum peak ΔF/F_0_ response from the animals in A. n=67-70; error bars are standard error of the mean. Mann-Whitney test; no significance. C. *Left:* Mean %ΔF/F_0_ response traces and *Right,* quantification of the maximum peak %ΔF/F_0_ responses to 30% sucrose stimulation of the labellum in the β’2 compartment of *MB301B*>*GCaMP6s::Brp::mCherry* flies fed a control (CD, *grey*) or high fat diet (FD, *rose*) n=31-32; shading and error bars are standard error of the mean. Mann-Whitney test; no significance. D. Latency-to-peak response times for the animals in C. n=31-32; error bars are standard error of the mean. Mann-Whitney test; no significance. E. *Left,* Mean %ΔF/F_0_ traces and *Right,* quantification of the maximum peak %ΔF/F_0_ responses to sucrose stimulation of the labellum in the β’2 compartment of *MB301B>GCaMP6s::Brp::mCherry* flies fed a control (CD, *charcoal*) and sugar diet (SD, *red*) supplemented with 75 µM OSMI-1. n=30-32; shading and error bars are standard error of the mean. Mann-Whitney test; no significance. F. The delay in the calcium responses quantified as latency in seconds (s) to maximum peak ΔF/F_0_ response from the animals in E. n=30-32; error bars are standard error of the mean. Mann-Whitney test; no significance.

To further probe this question, we fed flies a high fat diet (FD), which has the same caloric content of the high sugar diet (SD) and promotes fat accumulation, but does not decrease the responses of the *Gr64f+* sensory neurons to sugar stimuli (May et al. 2019). If changes in PAM-β’2 responses to sugar taste occur because of the metabolic side-effects of high nutrient density (i.e, fat accumulation) – rather than via changes in the sweet sensory neurons output – we would expect a FD to also induce PAM-β’2 dysfunction. However, a FD diet had no effect on the PAM-β’2 responses to sucrose or water stimulation of the labellum in *MB301B*>*GCaMP6s::Bruchpilot::mCherry* flies (Figure 3C, D and Supplemental Figure 2A, B). Together, these two lines of evidence argue that the dysfunction in the processing of sweet taste stimuli in the PAM-β’2 neurons of flies on a SD is linked to alterations in the peripheral sensory processing of sugar taste caused by high dietary sugar.

To test this hypothesis directly, we examined the effect of correcting sweet taste sensation on the responses of the PAM-β’2 *MB301B* neurons to sugar. To rescue the sweet taste deficits caused by a high sugar diet we fed flies an inhibitor of the metabolic-signalling enzyme O-GlcNAc-Transferase (OGT), which we previously found to be responsible for decreasing sweet taste on a SD (May et al. 2019). In accordance with our previous findings on OGT (May et al. 2019), supplementing the flies’ diet with 75 µM of OSMI-1 (OGT-small molecule inhibitor 1) resulted in no changes in PER between a CD and SD (Supplemental Figure 2C). In these flies, the calcium responses of PAM-β’2 neurons to sucrose stimulation of the labellum were identical in SD+OSMI and CD+OSMI flies, consistent with the idea that deficits in the peripheral responses drive impairments in the central processing of sweetness (Figure 3E, F).

Together, these orthogonal lines of evidence show that the impairments in the central processing of sweet sensory information by DANs are mediated by deficits in peripheral sweet taste responses.

### Correcting the activity of PAM DANs rescues feeding behavior

We previously showed that a diet-dependent dulling of sweet taste drives higher feeding behavior and weight gain by increasing the size and duration of meals (May et al. 2019). Since sweet taste deficits underlie the changes in PAM-β’2 activity, we reasoned that impairments in the central processing of orosensory signals may also play a role in promoting higher feeding in animals fed a high sugar diet. Specifically, if PAM-β’2 neurons were critical for integrating sweet taste information into feeding decisions, then correcting their activity may also prevent increased eating and weight gain when flies are exposed to a SD. To test this possibility we expressed the light-activated cation channel *ReaChR* (Inagaki et al. 2014) in the *MB301B* neurons, and used the optoFLIC, a feeding frequency assay (Ro, Harvanek, and Pletcher 2014) modified for closed-loop optogenetic stimulation (May et al. 2019), to stimulate the activity of PAM-β’2 neurons only when the flies were interacting with the food starting at day 3. *MB301B*>*ReaChR* flies that did not receive retinal supplementation (ATR, all-*trans*-retinal is required to form a functional light-sensitive opsin) exhibited the characteristic increase in feeding behavior on 20% sucrose (Figure 4A, *rose line*); however, *MB301B*>*ReaChR* +ATR animals, which were activated by light, had stable feeding for 10 days (Figure 4A, *peach line*). Control animals on 20% sucrose had more feeding interactions per meal and longer meal duration with more days on the SD (Figure 4B and C, *rose lines*), consistent with our previous data (May et al. 2019). In particular, we found that a SD induced a lengthening of the peak-to-end of the meal by ∼4 hours, suggesting that the satiation process is delayed in these animals (Figure 4D, *rose line*). However, feeding-paired stimulation of PAM-β’2 neurons stabilized the size and duration of the meal, as well as the time to satiation, over the entire duration of the experiment (Figure 4B, C, and D, *peach lines*). Interestingly, activation of the *Gr64f+* sweet taste neurons also corrected these two aspects of meal structure (May et al. 2019). Importantly, flies in which these PAM-β’2 DANs were activated still developed taste deficits on a SD (Supplemental Figure 3A), arguing against the possibility that PAM-β’2 DANs activation prevents increased feeding by rescuing the taste changes in the *Gr64f+* neurons. Instead, our data suggest that PAM-β’2 DANs modulate meal structure and feeding behavior by integrating the sensory signal from the periphery. In accordance with the stable feeding patterns recorded on the optoFLIC, we found that activation of PAM-β’2 DANs also prevented diet-induced obesity in animals fed high dietary sugar (Supplemental Figure 3B). Interestingly, PAM-β’2 DANs labeled by *MB301B* seem to play a unique role in this process. Activation of different subpopulations of PAM-β’2 with 8 distinct *GAL4* transgenes (Aso and Rubin 2016) (*MB056B*, *MB109B*, *MB042B*, *MB032B*, *MB312B*, *MB196B*, *MB316B*, some of these also express in γ4) failed to rescue diet-induced obesity (Supplemental Figure 3C). Further, flies with activation of nutrient-responsive PAM DANs (Yamagata et al. 2015; Huetteroth et al. 2015), which express in β2, still accumulated fat as controls when fed high dietary sugar, suggesting that effects of *MB301B* neuron activation come from the β’2 compartment alone (Supplemental Figure 3D).

**Figure 4:**
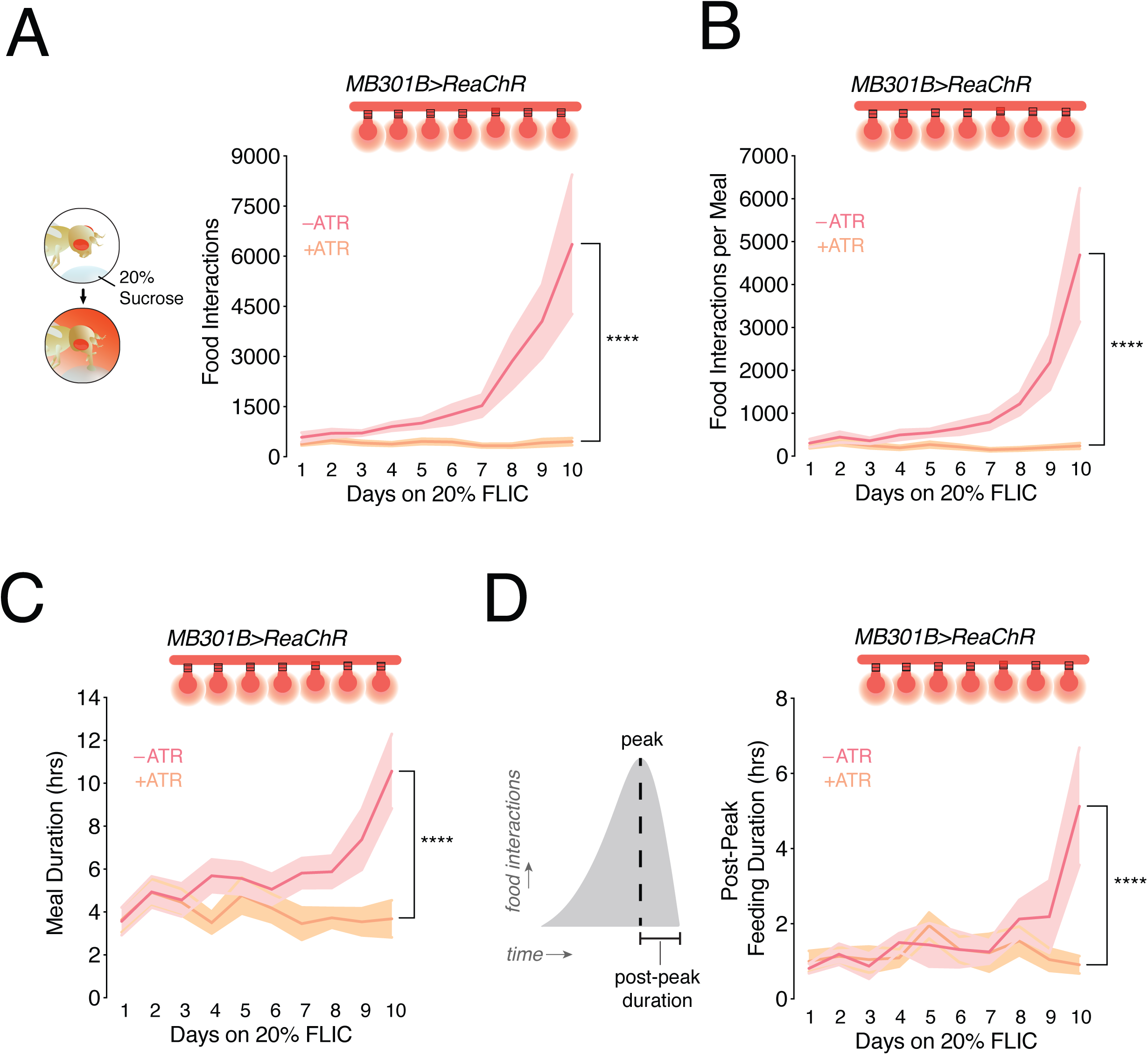
Closed-loop optogenetic activation of PAM-β’2 neurons corrects meal size and duration in flies fed a high sugar diet. A. *Left:* Conceptual schematic for the closed-loop optogenetic FLIC (optoFLIC), wherein a fly feeding on the 20% sucrose food triggers delivery of the red light during the food interaction. *Right:* Mean number of food interactions per day for *MB301B>ReaChR* flies fed 20% sucrose on the optoFLIC. Closed-loop light delivery was started on day 3 (indicated with *red light bulbs*). Control flies were not fed retinal (-ATR, *rose*), while experimental animals were fed retinal food before starting the experiment on the optoFLIC (+ATR, *peach*). n=8-11; shading is standard error of the mean. Two-way Repeated Measure (RM) ANOVA; \*\*\*\**p*<0.0001, Time by Retinal-treatment interaction. B. The size of the evening meal measured as the number of food interactions per meal for animals in A. n=8-11; shading is standard error of the mean. Two-way RM ANOVA; \*\*\*\**p*<0.0001, Time by Retinal-treatment interaction. C. The duration of the evening meal for animals in A. n=8-11; shading is standard error of the mean. Two-way RM ANOVA; \*\*\*\**p*<0.0001, Time by Retinal-treatment interaction. D. *Left*, schematic of an evening meal, and *Right*, mean duration of the portion of the evening meal after the peak (satiation) in animals from A. n=8-11; shading is standard error of the mean. Two-way RM ANOVA; \*\*\*\**p*<0.0001, Time by Retinal-treatment interaction.

### PAM-β’2 activity modulates the feeding rate during a meal

Since the FLIC records feeding interactions every 200 milliseconds (Ro, Harvanek, and Pletcher 2014), we used this information to look at how feeding rate changed during a meal, as this has been linked to the process of satiation. To do this, we first calculated the number of feeding events per meal, where a feeding event is defined as a succession of consecutive feeding interactions above an established signal threshold, (see Methods, and (Ro, Harvanek, and Pletcher 2014)). We next divided the number of events per meal by the duration of each meal per day to obtain a feeding rate and to control for the fact that meals last longer on a SD. We found that both the feeding events per meal and the feeding rate increased with chronic exposures to high dietary sugar (Figure 5A and B). However, optogenetic stimulation of PAM-β’2 prevented these increases and maintained a stable number of events and a constant feeding rate per meal over the duration of the experiment. We next examined whether the feeding rate changed during the course of the meal, by calculating it *before* and *after* the peak of meal feeding (Figure 5C, diagram). The feeding rate past the peak of the meal increased with time in animals fed 20% sucrose, but stayed the same in flies with activation of PAM-β’2 neurons (Figure 5C). Interestingly, the pre-peak eating also increased gradually with exposure to high dietary sugar (Figure 5D). Together, these data suggest that diet-dependent impairments in PAM-β’2 neurons promote overfeeding by impairing satiation, and specifically by affecting the feeding rate during a meal. Since, PAM-β’2 neurons process sensory experiences from the periphery, our experiments argue that this phenomenon is connected to sensory-enhanced satiation. Together we propose that the central processing of sensory experiences during a meal by PAM-β’2 DANs, controls feeding rate and sensory-enhanced satiation. This process is altered by high dietary sugar, leading to an attenuated satiation process and higher feeding (Figure 5E).

**Figure 5:**
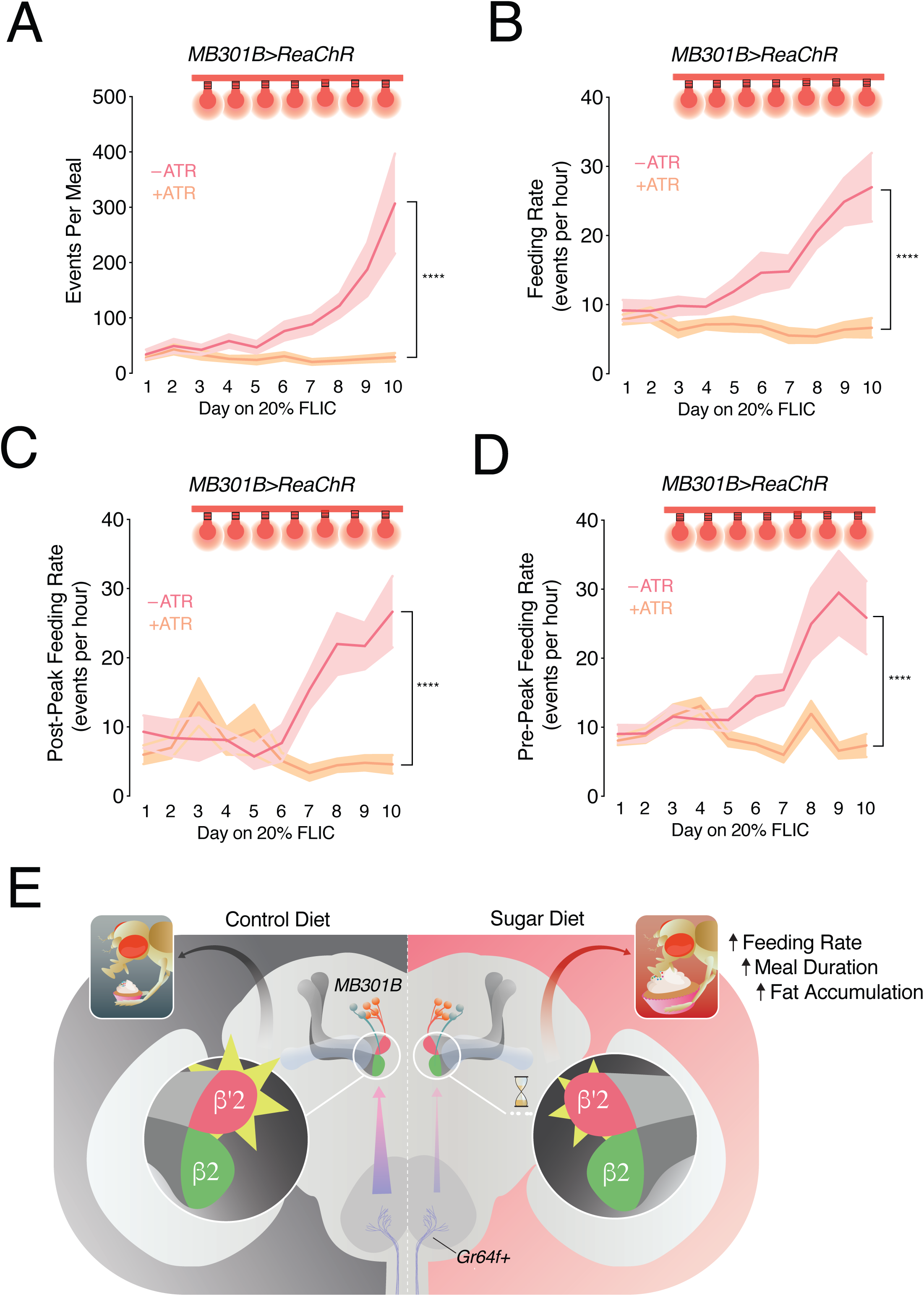
Feeding rate is modulated by a high sugar diet and controlled by the activity of PAM-β’2 neurons. A. The mean of total feeding events per meal in *MB301B>ReaChR* flies with (- ATR, *rose*) or without (+ATR, *peach*) retinal pretreatment. A feeding event is calculated as the number of consecutive licks above and below the signal threshold (see Methods). n=8-11; shading is standard error of the mean. Two-way Repeated Measures (RM) ANOVA; \*\*\*\**p*<0.0001, Time by Retinal-treatment interaction. B. The feeding rate per meal, calculated as the mean number of events per hour of mealtime in the animals from A. n=8-11; shading is standard error of the mean. Two-way RM ANOVA; \*\*\*\**p*<0.0001, Time by Retinal-treatment interaction. C. Quantification of the mean feeding rate *after* the peak of the meal in animals from A. n=8-11; shading is standard error of the mean. Two-way RM ANOVA; \*\*\*\**p*<0.0001, Time by Retinal-treatment interaction. D. Quantification of the mean feeding rate *before and including* the peak of the meal from flies in A. n=8-11; shading is standard error of the mean. Two-way RM ANOVA; \*\*\*\**p*<0.0001, Time by Retinal-treatment interaction. E. Model of the sweet taste and PAM DAN circuit changes when flies are fed a control (*left*) or high sugar diet (*right*): a decrease in the output of the *Gr64f+* neurons (*lavender axons, arrows*) contributes to a decrease (*yellow rays*) and a delay (*hourglass*) in the central processing of sweet taste information in the *PAM-β’2* terminals (*rose*), promoting higher feeding.

## DISCUSSION

In this study we found that diet-dependent changes in sensory perception promote feeding and weight gain by impairing the central dopaminergic processing of sweet taste information. When animals consume a high sugar diet, the responses to sweet taste of a distinct population of PAM DANs innervating the β’2 compartment of the MB are decreased and delayed. These alterations in dopaminergic processing increase the eating rate and extend the duration of meals, leading to attenuated satiation, higher feeding and weight gain (Figure 5E). Interestingly, we observed a reduction in PAM DAN responses only when flies ate diets that resulted in sweet taste deficits; consumption of an equal calorically-rich lard diet that did not impact taste had no effect on the PAM DANs responses. Similarly, animals fed high dietary sugar exhibited differences in PAM-β’2 responses to sweet, but not water taste stimuli, reinforcing the idea that PAM DAN alterations occur because of lower signal transmission from the sensory neurons (Figure 5E). Indeed, correcting sweet taste deficits also prevented impairments in PAM-β’2 responses. Thus, we propose a model where diet-dependent changes in taste intensity and sensitivity reduce the central processing of sensory stimuli to cause weaker and attenuated satiation. A weakness of the current study is that we were unable to follow the transmission of the taste signal from the primary sensory neurons through the different circuits that eventually communicate with PAM. Studies that will identify taste projection neurons genetically will allow us to further probe this point in the future.

Studies in rodents and humans have delineated the importance of sensory signals to modulate satiation and terminate meals. This process, termed sensory-enhanced satiation (Chambers, McCrickerd, and Yeomans 2015), plays an early role in the satiety cascade before post-oral nutrient-derived signals consolidate satiety (J. E. Blundell, Rogers, and Hill 1987; Bellisle and Blundell 2013). Studies show that higher sensory intensity and oral exposure promote stronger satiation (Bolhuis et al. 2011; Ramaekers et al. 2014). For example, high sensory characteristics, such as saltiness and sweetness, enhanced the satiating effect of both low and high energy test drinks (Yeomans and Chambers 2011; Yeomans et al. 2014), decreased consumption of pasta sauce (Yeomans 1998, 1996), yoghurt (Vickers, Holton, and Wang 2001) and tea (Vickers and Holton 1998). However, the neural basis for this phenomenon is unknown. Here we characterized the circuit-based mechanisms of sensory-enhanced satiation by exploiting the simplicity of the fruit fly system. We show that sensory-enhanced satiation involves the central dopaminergic processing of peripheral sweet taste stimuli by a dedicated group of PAM-β’2 neurons. Given the role of PAM DANs transmission in reinforcing appetitive memories (Burke et al. 2012; Liu et al. 2012), this discovery is significant because it suggests that satiation may involve a learning or rewarding component and that diet composition may direct food intake by influencing this aspect. Indeed, sensory cues function as a predictor of nutrient density and set expectations for how filling different types of foods should be (Chambers, McCrickerd, and Yeomans 2015; McCrickerd and Forde 2016; Yeomans 2017). This information could be used to modulate the feeding rate during the meal and initiate the process of meal termination without relying uniquely on nutrient-derived cues, which arrive later (Bellisle and Blundell 2013; J. E. Blundell, Rogers, and Hill 1987). The idea that sensory cues could set cognitive expectations about the fullness of future meals is also in line with the known roles of DA in promoting the formation of appetitive memories. In fruit flies, PAM DANs promote the formation of short-term associative memories based on taste and long-term associative memories based on nutrient density by modulating plasticity of the postsynaptic Mushroom Body Output Neurons (MBONs) (Cohn, Morantte, and Ruta 2015; Owald et al. 2015). MBONs are, in turn, connected to pre-motor areas like the Central Complex (Aso et al. 2014) – the fly genetic and functional analog of the basal ganglia (Strausfeld and Hirth 2013) – providing an anatomical route to modulate aspects of feeding such as proboscis extension (Chia and Scott 2019), the analogue of licking or chewing rate. Interestingly, some MBONs receive input from both the taste (β’2) and nutrient (γ5) compartments, raising the possibility that sensory and nutrient memories may be integrated in the same cells to regulate different aspects of the satiety cascade (satiation vs. satiety). In flies, the mode and timing of DA delivery onto the MBONs is critical to establish the strength and valence of the associations (Handler et al. 2019). The delay and decrease we measured in animals on a high sugar diet could impair MBON synaptic plasticity and the formation of new appetitive memories (Cohn, Morantte, and Ruta 2015; Owald et al. 2015). If this is the case, we would expect that flies on this diet may be insensitive to new learning, use old food memories to predict the filling effects of the meal, and thus overshoot their food intake. This is consistent with the idea elegantly espoused by (Kroemer and Small 2016) who explain the decrease in DA transmission with diet or obesity in a reinforcement learning framework. A different possibility, however, is that alterations in PAM DAN processing are not related to reinforcement learning per se, but instead to a decrease in overall reward receipt. In this light, sensory signals would cue reward not learning, and the pleasure experienced during eating would promote satiation and curb food intake. The idea that decreases in the sensitivity of the reward system increases food intake has been described as the “reward deficit” theory of obesity (Wang, Volkow, and Fowler 2002), which also draws a parallel between the effects of drugs of abuse and that of sugar on the brain. Our results are consistent with both reinforcement learning and reward deficit scenarios, as well as with other integrated theories of obesity (Stice and Yokum 2016); future experiments examining the role of circuits downstream of PAMs, and especially the involvement of MBONs, will differentiate between these possibilities. Our study also adds to the current body of evidence connecting diet with DA alterations in mammals (Geiger et al. 2009; van de Giessen et al. 2013; Kroemer and Small 2016; Friend et al. 2017; DiFeliceantonio and Small 2019). In particular, we speculate that some of the changes in DA transmission observed with diet exposure in rodents and humans may be due to impairments in sensory processing, since humans and rodents also process the taste and nutritive properties of sugar separately (Tellez et al. 2016; Thanarajah et al. 2019).

In conclusion, our experiments demonstrate that by reducing peripheral taste sensation, a high sugar diet impairs the central DA processing of sensory signals and weakens satiation. These studies forge a causal link between sugar – a key component of processed foods – taste sensation, and weakened satiation, consistent with the fact that humans consume more calories when their diets consist of processed foods (Hall et al. 2019). Given the importance of sensory changes in initiating this cascade of circuit dysfunction, understanding how diet composition mechanistically affects taste is imperative to understand how the food environment directs feeding behavior and metabolic disease.

## METHODS

### Fly Lines and Preparation

All flies were maintained at 25°C in a humidity-controlled incubator with a 12:12 hours light/dark cycle. For all experiments, males were collected under CO_2_ anesthesia, 2-4 days following eclosion, and housed in groups of 20-30 within culture vials. The *GAL4*/*UAS* system was used for cell-type specific expression of transgenes. Stocks used are listed in the table below; *w^1118^Canton-S* was used as control.

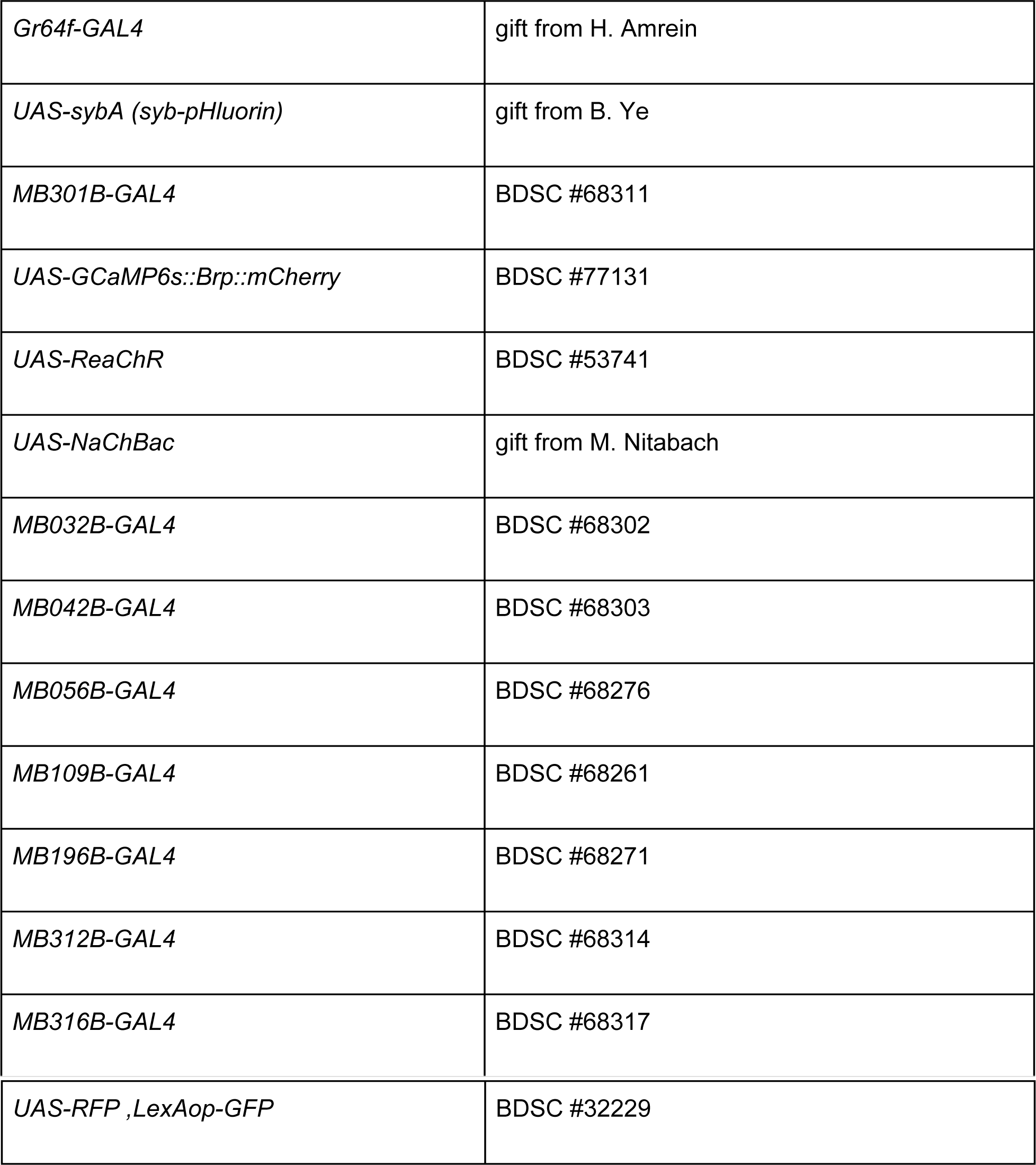

|  |  |
| --- | --- |
| *Gr64f-GAL4* | gift from H. Amrein |
| *UAS-sybA (syb-pHluorin)* | gift from B. Ye |
| *MB301B-GAL4* | BDSC #68311 |
| *UAS-GCaMP6s::Brp::mCherry* | BDSC #77131 |
| *UAS-ReaChR* | BDSC #53741 |
| *UAS-NaChBac* | gift from M. Nitabach |
| *MB032B-GAL4* | BDSC #68302 |
| *MB042B-GAL4* | BDSC #68303 |
| *MB056B-GAL4* | BDSC #68276 |
| *MB109B-GAL4* | BDSC #68261 |
| *MB196B-GAL4* | BDSC #68271 |
| *MB312B-GAL4* | BDSC #68314 |
| *MB316B-GAL4* | BDSC #68317 |
| *UAS-RFP ,LexAop-GFP* | BDSC #32229 |

### Dietary Manipulations

Flies were transferred to each diet 2-4 days after eclosion in groups of 30 animals per vial and fed on experimental diets (SD or FD) for 7 days with age-matched controls on CD.

The composition and caloric amount of each diet was as below:

- “Control Diet/CD” was a standard cornmeal food (Bloomington Food B recipe), with approx. 0.6 cal/g.
- “Sugar Diet/SD” was 30 g of table sugar added to 89 g Control Diet for 100 mL final volume of 30% sugar w/v, with approx. 1.4 cal/g.
- “Fat Diet/FD” was 10 mL of melted lard added to 90 mL of liquid Control Diet for 100 mL final volume of 10% lard v/v, with approx 1.4 cal/g.
- For diets supplemented with OSMI-1, the inhibitor was dissolved in 55% DMSO for a stock concentration of 500 µM, and then diluted 3:20 in liquid Control or Sugar Diet for a final concentration of 75 µM in food.
- For diets supplemented with all-*trans*-retinal, retinal was dissolved in 95% EtOH for a stock concentration of 20 mM, then diluted 1:100 in liquid Control Diet for a final concentration of 200 µM in food.
- Diets on the FLIC were 5% and 20% w/v D-sucrose in 4 mg/L MgCl_2_.

### In vivo Imaging

Adult age-matched male flies, following 7 days of CD or SD, were fasted on a wet Kimwipe for 18-24 hours before prepping for *in vivo* confocal laser imaging. As previously described (May et al. 2019; LeDue et al. 2015), the preparation consisted of a fly affixed to a 3D-printed slide with melted wax around the head and on the dorsal part of the thorax. Distal tarsal segments were removed prevent interference of the proboscis stimulus, and the proboscis was wax-fixed fully extended with the labellum functional and clear of wax so that proboscis contraction and extension could not perturb the brain’s position. A glass coverslip was placed such that artificial hemolymph (108 mM NaCl, 8.2 mM MgCl_2_, 4 mM NaHCO_3_, 1 mM NaH_2_PO_4_, 2 mM CaCl_2_, 5 mM KCl, 5 mM HEPES) placed over the head did not touch the proboscis. Data were acquired with a FV1200 Olympus confocal microscope, a 20x water immersion objective, and a rate of 0.254 s per frame. Stimuli consisted of a brief touch of a small Kimwipe soaked in milliQ water or 30% sucrose solution to the labellum.

### Optogenetic Stimulation for Fly-to-Liquid-food Interaction Counter (optoFLIC)

optoFLIC was run as previously described (May et al. 2019). Briefly, adult flies 3-5 days past eclosion were placed on ATR food and kept in the dark for 3 days until starting the optoFLIC. optoFLIC experiments were run in an incubator with consistent 25°C and 30-40% humidity, on a dark/dark light cycle to prevent ambient-light activation of the ReaChR. Following two days recording of feeding activity on the FLIC food without LED activation, a protocol for closed-loop feeding-triggered LED activation was begun. The LED activation protocols were as follows: For experiments with MB301B>ReaChR, 200 ms of red (∼627 nm) light pulsing at frequency 60 Hz and with a pulse width of 4 ms was triggered by every food interaction signal over 10.

### Immunofluorescence Staining

Immunofluorescence protocol was performed as described in (Dus et al. 2015). Briefly, brains were dissected in 1xPBS from male *MB301B>RFP* flies 3-5 days post-eclosion, then fixed in 4% paraformaldehyde in 1xPBS for 20 min, blocked in blocking buffer (10% normal goat serum, 2% Triton X-100 in 1xPBS), and incubated overnight at RT in anti-TH (rabbit polyclonal Ab from Novus Bio) 1:250 in dilution buffer (1% normal goat serum, 0.25 Triton X-100 in 1xPBS). Secondary antibody was goat anti-rabbit Alexa Fluor 488 diluted 1:500 in dilution buffer, and brains were washed then incubated with secondary antibody overnight at RT. Brains were mounted in FocusClear between two coverslips and imaged within 24 hours.

### Triacylglyceride (TAG) Assay

Following the protocol in (Tennessen et al. 2014), we assayed total TAG levels normalized to total protein in whole male flies. To assay, flies were CO_2_-anesthetized and flash frozen. Pairs of flies were homogenized in lysis buffer (140 mM NaCl, 50 mM Tris-HCl pH 7.4, 0.1% Triton-X) containing protease inhibitor (Thermo Scientific). Separation by centrifugation produced a supernatant containing total protein and TAGs. Protein reagent (Thermo Scientific Pierce^TM^ BCA Protein assay) was added to the supernatant and the standards and incubated for 30 min at 37°C, then tested for absorbance at 562 nm on a Tecan Plate Reader Infinite 200. TAG reagent (Stanbio Triglycerides LiquiColor Test) was added to supernatant and standards, incubated for 5 min at 37°C, then tested for absorbance at 500 nm.

### Proboscis Extension Response

Flies were fasted for 24h in a vial with a Kimwipe dampened with 2 mL of milliQ-filtered deionized (milliQ DI) water and tested for the proboscis extension response (PER) (Shiraiwa and Carlson 2007). Water and all tastants were tested manually via a solution-soaked Kimwipe. Sucrose solutions were dissolved in milliQ water and presented in descending order by concentration. Groups of 10-15 flies were tested simultaneously.

### Imaging Data Analysis

For each fly, ΔF/F_0_ was calculated from a baseline of 10 samples recorded just prior to the stimulus (sucrose or water). Area under the curve (AUC) was calculated by summing the ΔF/F_0_ values from the initiation of the response to its end. Peak ΔF/F_0_ is the single maximum acquired within a response, and latency to peak was calculated by determining the time between the stimulus delivery and the peak response.

### optoFLIC Data Analysis

OptoFLIC analysis of daily food interactions, meal size, and meal duration was performed as previously described (May et al. 2019). R code used can be found on Github (github.com/ chrismayumich/May_et_al_optoFLIC). Briefly, food interactions were determined by calculating a moving baseline on the raw data and selecting signals which surpassed threshold above baseline. These signals were then summed in 30-minute bins. From the binned data, daily food interactions and the start and end of meals were calculated. The evening meal was used for all meal-based calculations to control for variability in meal shape. Meal size and duration were derived using meal start and end. Post-peak feeding duration was quantified as *[(time of meal end) - (time of meal peak)]*.

An event is defined as a string of consecutive food interactions. R code used to extract event information can also be found on Github. To calculate events per meal, the number of events between the meal start and meal end per meal were summed for each fly. Feeding rate was quantified as *[(events per meal) / (meal duration)]* per meal per fly. Pre- and post-peak feeding rates were quantified, using the time of the meal peak determined by food interactions, also used to calculate post-peak feeding duration, as *[(number of events pre- or post-peak) / (pre- or post-peak feeding duration)]*. Pre-peak feeding duration was quantified as *[(time of meal peak) - (time of meal start)]*.

## AUTHOR CONTRIBUTIONS

CEM conducted all the experiments, with the exception of PER. MD carried out the PER experiments and supervised the project. JR, JG, and ED helped with the TAG measurements. MD and CEM wrote the manuscript together.

## ACKNOWLEDGEMENTS

We thank Carrie Ferrario, Shelly Flagel, and Josie Clowney for helpful discussions and Scott Pletcher for his continuous assistance with the FLIC. We also thank all the researchers that shared protocols and fly lines (listed in the Methods). Julia Kuhl designed some of the graphics for the manuscript. This work was funded by NIH R00 DK-97141 and NIH 1DP2DK-113750, a NARSAD Young Investigator Award, the Klingenstein-Simons Fellowship in the Neurosciences, and the Rita Allen Foundation (to M.D.), and by Training Grant T32-GM008322 (to C.E.M.)

## DECLARATION OF INTEREST

The authors declare no competing interests.

---

**Supplemental Figure 1 (related to Figure 2):**
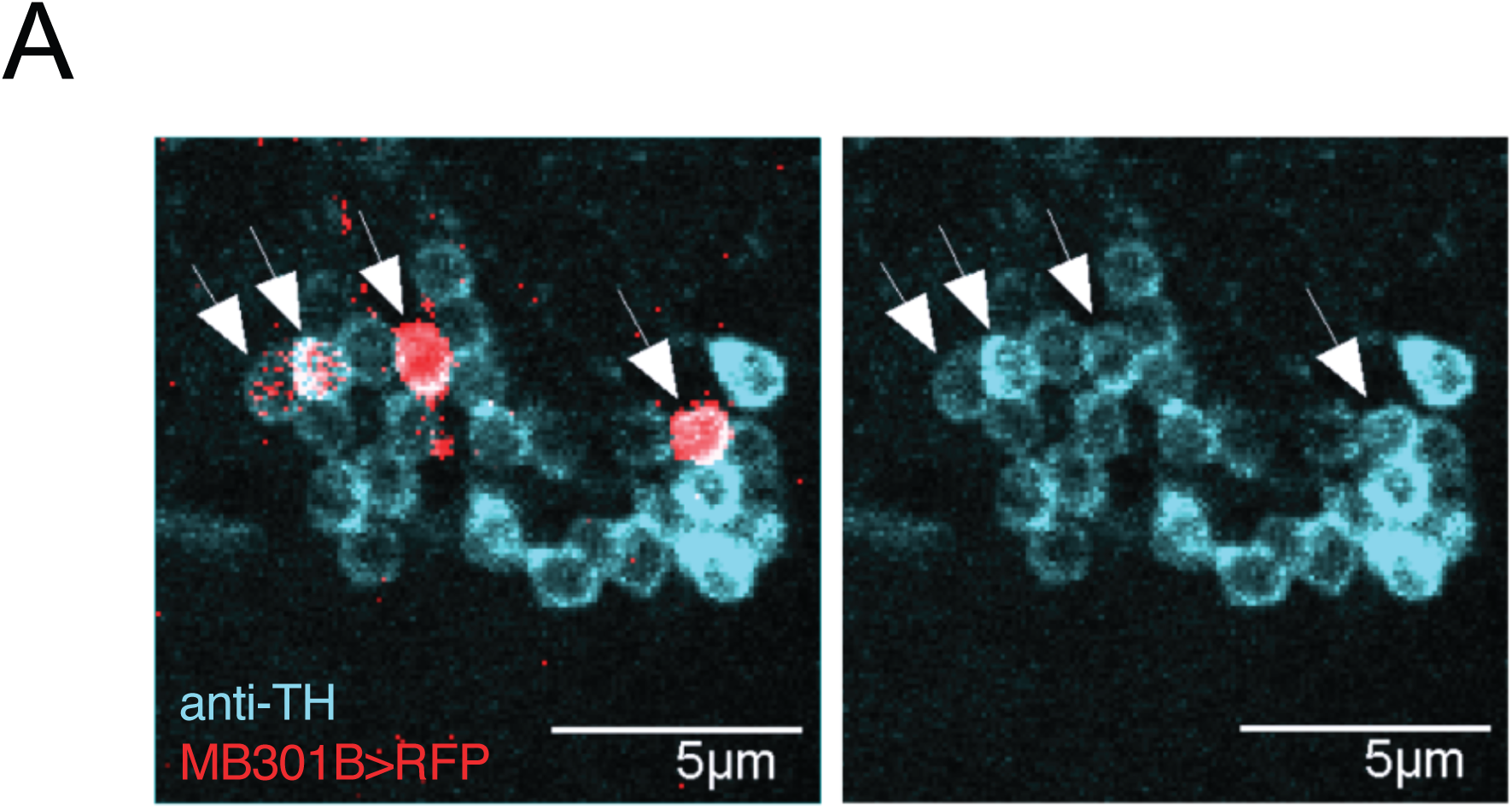
A. Confocal fluorescence image of the PAM cluster neurons stained with an antibody against tyrosine hydroxylase (TH, *cyan*) in *MB301B>RFP* flies. Arrows indicate *MB301B* cell bodies (*red*) which are also positive for TH. Scale bar = 5 µm.

**Supplemental Figure 2 (related to Figure 3):**
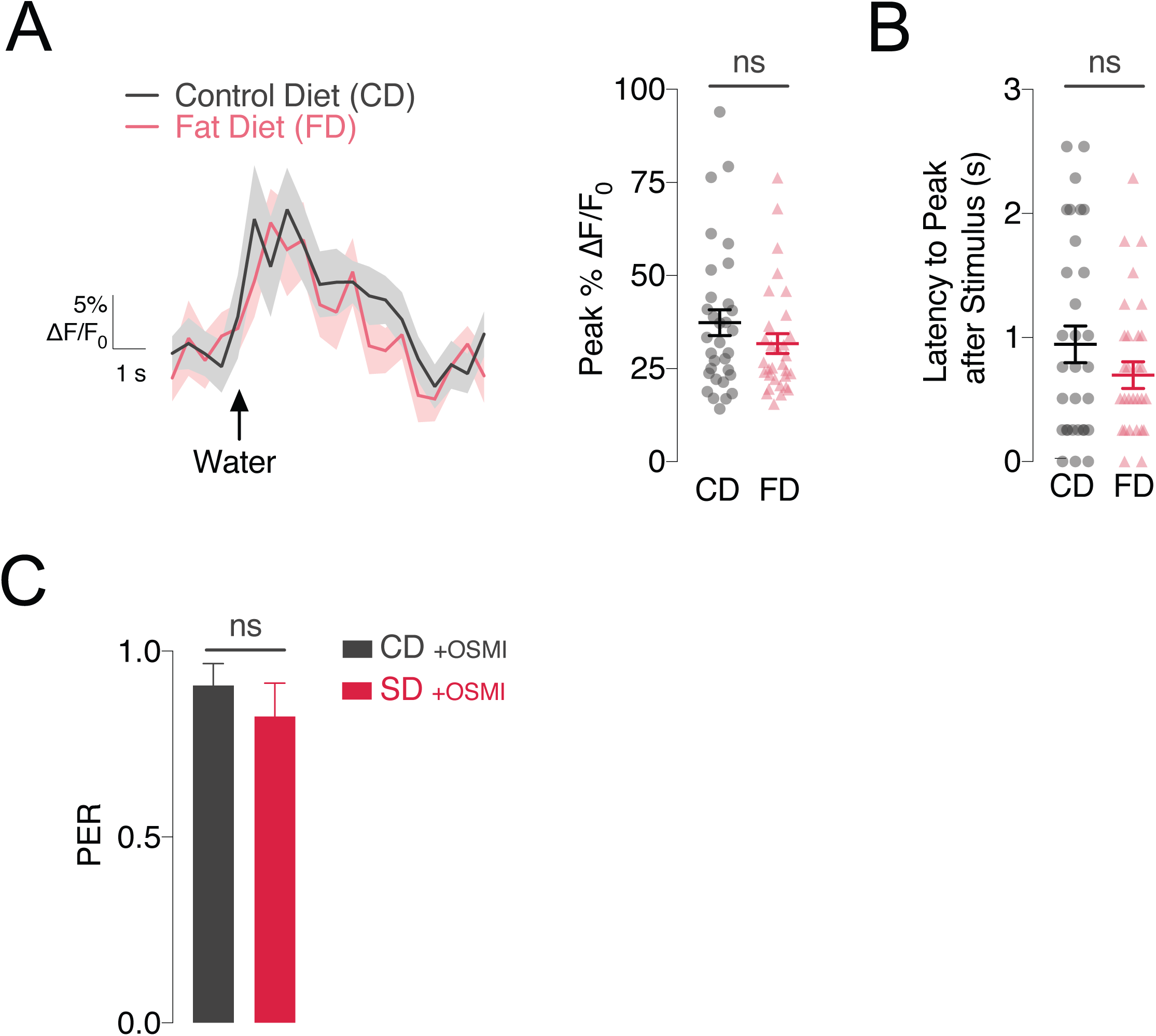
A. *Left,* Mean %ΔF/F_0_ traces and *Right,* quantification of the maximum peak %ΔF/F_0_ responses to stimulation of the labellum with water in the β’2 compartment of *MB301B>GCaMP6s::Brp::mCherry* flies fed a control (CD, *grey*) and high fat diet (FD, *rose*). n=31-32; shading and error bars are standard error of the mean. Mann-Whitney test; no significance. B. Latency-to-peak response times for the animals in A. n=31-32; error bars are standard error of the mean. Mann-Whitney test; no significance. C. Mean score of Proboscis Extension Response (PER) to stimulation of the labellum with 30% sucrose in male *w^1118^CS* flies, fed a control (CD, *charcoal*) and high sugar (SD, *red*), supplemented with 75 µM OSMI-1 (OSMI-1). n=18 per group; error bars are standard error of the mean. Mann-Whitney test; no significance.

**Supplemental Figure 3 (related to Figure 4):**
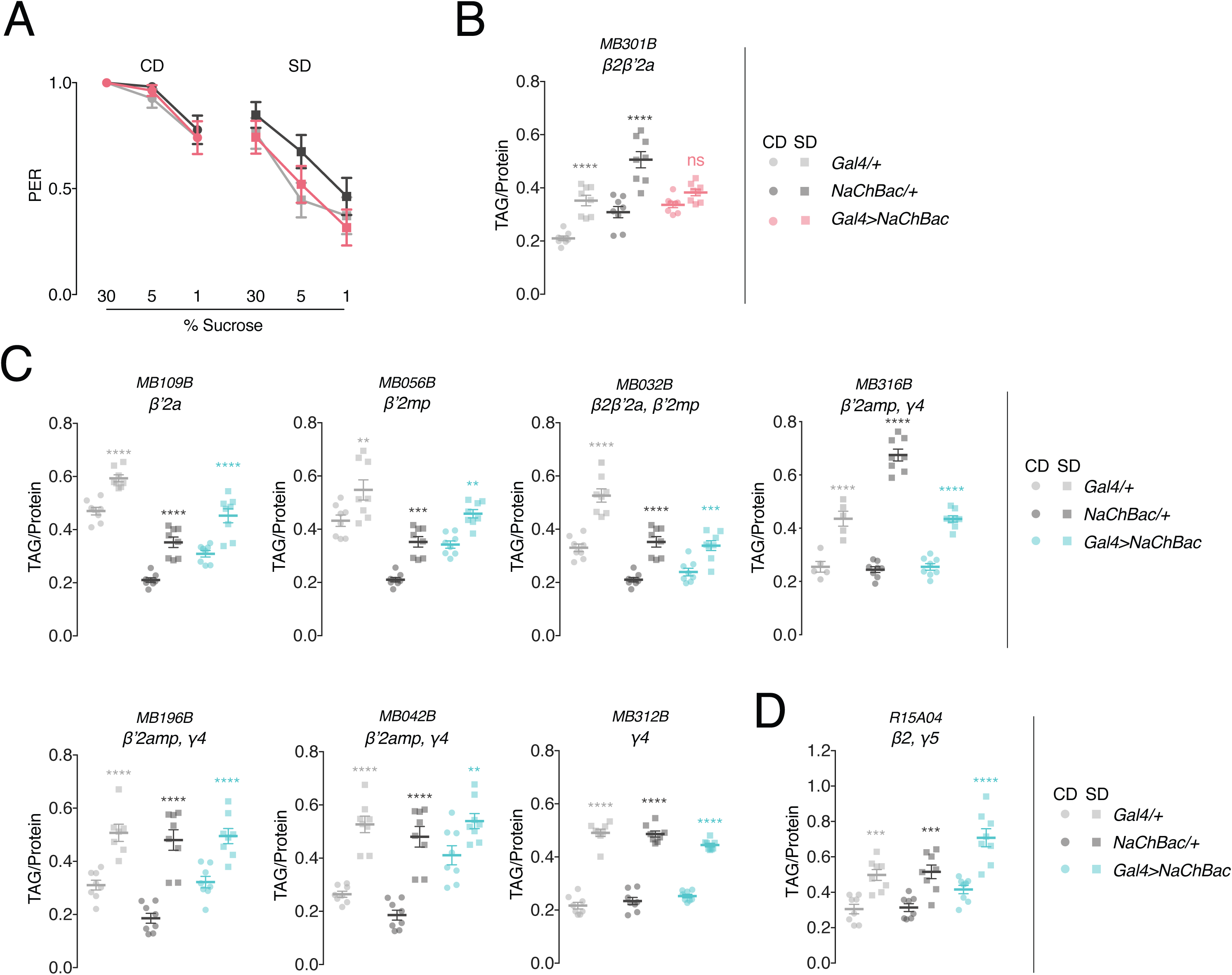
A. Mean proboscis extension response (PER) to sucrose stimulation of the labellum (30%, 5%, and 1%) in *MB301B>NaChBac* flies following 7 days of exposure to a control (CD, *left*) or sugar diet (SD, *right*). n=27 per condition; error bars are standard error of the mean. Kruskal-Wallis with Dunn’s multiple comparisons, no significance. B. Mean triacylglyceride (TAG) content normalized to protein of *MB301B>NaChBac* flies and single transgenic control male flies fed a CD or SD for 7 days. n=8 per condition; error bars are standard error of the mean. Two-way ANOVA with Sidak’s multiple comparisons test; \*\*\*\**p*<0.0001, comparison to CD within genotype. C. Mean TAG content normalized to protein of male flies with expression of *UAS-NaChBac* in different subsets of PAM neurons innervating β’2 or γ4 regions of the mushroom body. n=5-8; error bars are standard error of the mean. Two-way ANOVA with Sidak’s multiple comparisons test; \*\**p*<0.01, \*\*\**p*<0.001, \*\*\*\**p*<0.0001, comparison to CD within genotype. Legends are on the right of the figure. D. Mean TAG content normalized to protein of male flies with expression of *UAS-NaChBac* in nutrient-reward PAM neurons which innervate the β2 compartment of the mushroom body. n=8 per condition; error bar is standard error of the mean. Two-way ANOVA with Sidak’s multiple comparisons test; no significance, comparison to CD within genotype.

## Notes

https://github.com/chrismayumich/May_et_al_optoFLIC/

